# Insulin Resistance Increases TNBC Aggressiveness and Brain Metastasis via Adipocyte-derived Exosomes

**DOI:** 10.1101/2024.05.01.592097

**Authors:** Yuhan Qiu, Andrew Chen, Rebecca Yu, Pablo Llevenes, Michael Seen, Naomi Y. Ko, Stefano Monti, Gerald V. Denis

## Abstract

Patients with triple negative breast cancer (TNBC) and comorbid Type 2 Diabetes (T2D), characterized by insulin resistance of adipose tissue, have higher risk of metastasis and shorter survival. Adipocytes are the main non-malignant cells of the breast tumor microenvironment (TME). However, adipocyte metabolism is usually ignored in oncology and mechanisms that couple T2D to TNBC outcomes are poorly understood. Here we hypothesized that exosomes, small vesicles secreted by TME breast adipocytes, drive epithelial-to-mesenchymal transition (EMT) and metastasis in TNBC via miRNAs. Exosomes were purified from conditioned media of 3T3-L1 mature adipocytes, either insulin-sensitive (IS) or insulin-resistant (IR). Murine 4T1 cells, a TNBC model, were treated with exosomes *in vitro* (72h). EMT, proliferation and angiogenesis were elevated in IR vs. control and IS. Brain metastases showed more mesenchymal morphology and EMT enrichment in the IR group. MiR-145a-3p is highly differentially expressed between IS and IR, and potentially regulates metastasis.

**Significance:** IR adipocyte exosomes modify TME, increase EMT and promote metastasis to distant organs, likely through miRNA pathways. We suggest metabolic diseases such as T2D reshape the TME, promoting metastasis and decreasing survival. Therefore, TNBC patients with T2D should be closely monitored for metastasis, with metabolic medications considered.

## Introduction

Triple-negative breast cancer (TNBC) accounts for 11-20% of all breast cancers (1), lacks expression of estrogen receptor (ER) and progesterone receptor (PR) and human epidermal growth factor 2 receptor (HER2) (2), and consequently is not amenable to receptor-targeted therapeutics. TNBC cells demonstrate greater transcriptional plasticity than other breast cancer cell types and easily induce epithelial-to-mesenchymal transition (EMT), a gene expression program associated with increased migration, invasion and earlier appearance of distant metastases (3–5). Patients with TNBC are reported to have shorter survival than other breast cancer types (6,7), and forty-five percent of advanced stage TNBC patients are reported to develop metastasis into brain or distant organs (8). Brain metastasis in TNBC patients is prevalent and associates with high mortality (9,10). These difficult clinical features of TNBC have prompted a major research effort to understand underlying mechanism and develop novel therapeutics. Attention has recently focused on the role of the breast tumor microenvironment (TME) as a critical factor in TNBC initiation and progression, in particular the role of metabolically dysregulated breast adipocytes found in patients with metabolic co-morbidities (11–13). Incident TNBC associates with metabolic disorders in studies across different ethnic groups and populations. For example, one retrospective study of invasive breast cancer showed obesity was present in 49.6% of TNBC patients, but only in 35.8% of non-TNBC counterparts (*P*=0.0098) (14). A meta-analysis of eleven epidemiology studies reported that women with obesity have a 20% greater risk of developing TNBC compared to women without obesity (15). Metabolic syndrome (16) and dyslipidemia (17) are particularly implicated as drivers of TNBC aggressiveness and mortality. Although about 25% of the adult obese population is considered metabolically healthy (18), much of the obese population exhibits metabolic complications, particularly insulin resistance (IR) and Type 2 diabetes (T2D). Here, we focus on novel TME crosstalk, originating with metabolically dysregulated adipocytes observed in patients with IR and T2D, to provoke TNBC aggressiveness and metastasis, which may suggest new avenues for clinical decision making and management of TNBC in patients with obesity.

It is long established that obesity is a risk factor for breast cancer in post-menopausal women (19–22), which has been linked to the increased aromatase activity of adipose tissue as a driver of estrogen production and thereby proliferation in ER-positive breast cancer (23,24). Obesity-driven T2D is also a risk factor for progression of breast cancer, particularly hormone receptor-negative breast cancer (25,26). We previously showed in the Black Women’s Health Study that duration of T2D ≥5 years positively associates with incidence and distant metastases of ER-negative cancer (27,28). However, these population-level observations have lacked mechanistic interpretation. Cell culture and animal models have been used to investigate inflammatory cytokines associated with adipocyte IR (29), including interleukin-6 (30); leptin, which is produced by mature adipocytes and elevated in obesity; adiponectin, which is reduced; and leptin-adiponectin ratio, as critical modifiers of breast tumor cell proliferation (31). We previously showed that IR or insulin sensitive (IS) status of mature primary adipocytes in co-culture with ER+ breast cancer cells is a decisive determinant of EMT (32). More aggressive phenotypes are provoked in the IR context compared to IS. Intriguingly, we identified adipocyte-origin exosomes as critical drivers of breast cancer EMT (32). Exosomes are extracellular vesicles ranging in diameter from 50 to 150 nm that encapsulate RNA, protein and lipid molecules, and are released by many cell types (33). Muller and coauthors (34) have shown that melanoma cells treated with adipocyte-origin exosomes upregulate fatty acid metabolism through exosomal delivery of metabolic enzymes, thereby increasing aggressiveness and linking obesity to melanoma progression. It has also been reported that hepatocyte-derived exosomes promote IS in early onset obese mice, but induce IR in chronically obese mice, not via insulin signaling, but through proinflammatory activation of macrophages (35). For breast cancer, we proved that exosomes are essential carriers of information from IR adipocytes that promote cancer cell EMT and aggressiveness (32). Other groups have shown that exosomes as well as other changes related to obesity are important to fuel cancer growth and increase aggressiveness. For example, Muller and coauthors were the first to show that adipocyte-derived exosomes increase fatty acid metabolism in melanoma cell lines (36,37). Brown and coauthors illustrated how obesity can exacerbate breast cancer development (38). These observations provided the rationale for the studies we report here, where we show that adipocyte-origin exosomes drive brain metastasis of TNBC in an animal model.

Despite these considerations, the standard of care in breast medical oncology does not yet fully consider how IR and T2D modify the breast TME in TNBC to stratify outcomes. The close association between obesity-driven T2D and breast cancer (39) identifies a need to study the biological bases of metastatic risk and how adipocyte metabolism and insulin resistance of the T2D/obese breast TME might differentially affect breast cancer metastasis and progression. The findings of this paper will call for the evaluation of metabolic status of patients in the context of TNBC, which will have far-reaching impact on the scope of TNBC research in oncology, immunology and discovery of potential biomarkers.

## Materials and Methods

### Cell Culture

Murine 4T1 cells as a TNBC model and murine 3T3-L1 pre-adipocytes to be differentiated into mature adipocytes were obtained from the American Type Culture Collection. The 4T1 cells and 3T3L1 cells were cultured in RPMI1640 and DMEM medium, respectively, supplemented with 10% fetal bovine serum (FBS), and 100 units/mL penicillin, 10 μg/mL streptomycin (Corning), and incubated at 37°C with 100% humidity and 5% CO_2_. Cell lines were tested for mycoplasma monthly using a Mycoplasma Detection Kit (Invivogen) and used within 10 passages.

### Animals

All animal experiments were performed in accordance with approved procedures of the Institutional Animal Care and Use Committee (IACUC) of Boston University Medical Center. Four-week-old female BALB-c/J mice were purchased from Jackson Laboratory and acclimated in the Boston University Animal Facility for a week. The 4T1 cells (20,000) were exposed to exosomes on Day 0 and allowed to proliferate to a total of 800,000 to 1 million by Day 3. Mice were then injected with 50,000 4T1 cells in the fourth mammary fat pad and were sacrificed two weeks after injection.

### Differentiation of 3T3-L1 pre-adipocytes and induction of insulin resistance

Two days after cells reached confluency, differentiation was induced by addition of 1 μM dexamethasone, 0.5 mM isobutylmethylxanthine, and 1.67 μM insulin in DMEM for 48 hours. The medium was then replaced with maintenance medium containing 0.41 μM insulin. Insulin resistance was then initiated in the mature adipocytes by addition of 1 nM recombinant murine tumor necrosis factor (TNF)-α (Abcam, ab9740) in maintenance medium for 24 hours.

### Exosome Extraction

Methods were as previously described [31]. Briefly, conditioned media (typically 15 mL) was centrifuged at 300 × *g* for 10 min in a 15 mL conical tube to remove cells and debris. The supernatant was then transferred to an Amicon Ultra-15 (Millipore-Sigma; REF-UFC910008) 100K centrifugal filter and centrifuged at 3,500 × *g* for 15 min to reach a final volume of 0.5 ml, then exosomes were purified by size exclusion chromatography on a qEV Original column, using an automatic fraction collector (AFC, serial number: V1-0395, IZON). The size distribution and concentration of exosomes were determined before each biological experiment using NanoSight NS300 system (Malvern Panalytical). The ratio of exosomes to 4T1 cells was 50,000:1 on Day 0 of exposure, based on NanoSight quantitation.

### EMT Array

Total RNA was isolated from cells using RNeasy Plus Mini Kit (74136, Qiagen). RNA was then used to prepare cDNA using QuantiTect Reverse Transcription Kit (Qiagen, 205313). Mouse RT2 Profiler™ PCR Array, including Epithelial to Mesenchymal Transition (EMT) (MAHS-090Z) arrays, were purchased from Qiagen. cDNA sample was mixed with RT2 SYBR Green ROX qPCR Mastermix (Qiagen, 330522) and PCR reactions were performed and results were analyzed using a 7500 Fast Real-Time PCR instrument. Fold change and Log fold change was then calculated and plotted with Prism software.

### Ingenuity pathway analysis

To predict disease and function, and downstream pathways, data were analyzed using Ingenuity Pathway Analysis, IPA (QIAGE Inc). Core analysis was conducted, then comparison analysis was performed on all the conditions of each experiment, including replicates.

### Immunohistochemistry

Antibodies used for immunohistochemistry (IHC) from mouse tissue sections were anti-Vimentin (Abcam, ab92547, 1:1000), anti-Ki67 (Abcam, 9701S,RRID:AB_331535, 1:1000), anti-Cd31 (Abcam, Ab281583, 1:4000). Formalin-fixed, paraffin-embedded (FFPE) slides from primary tumor were deparaffinized and then placed in antigen retrieval buffer for 15 mins at 95°C. After slides reached room temperature and were washed with running water, endogenous peroxidase activity was blocked using 3% H_2_O_2_ solution for 10mins, followed by general blocking using 2% FBS in PBS for 60 mins. Slides were then incubated with primary antibody in humidified chamber at 4°C overnight. After incubation with secondary antibody at room temperature for 30 min, DAB Substrate Kit, Peroxidase (SK-4100, Vector Lab) were used per manufacturer’s instructions, followed by counterstain with hematoxylin for 15 secs. Dehydration was performed and semi-dried slides were covered by mounting media (Permount, Fisher Chemical) and coverslips.

### Migration and Invasion Assays

Cells were switched from normal medium to serum-free medium for 3 hours and subsequently plated in 24-well, 8-m pore size transwell plates (Thermo Fisher Scientific). The cells were plated in the upper well of the transwell inserts with serum-free media, and the bottom well was filled with RPMI media supplemented with 10% FBS to serve as a chemoattractant. Cells that remained on the upper side of the membrane 20 hours later were removed, whereas cells that had migrated were fixed with ice-cold methanol for 5 min at −20°C. Fixed cells were then stained with 1% crystal violet (v/v) in 2% ethanol for 15 min at room temperature. Images were captured by an EVOS XL Core digital inverted microscope. The percentage of migration was quantified using ImageJ software [National Institutes of Health (NIH), Bethesda, MD] and then converted to relative percent migration/invasion by comparing each condition to the control condition.

### Morphology Analysis

Images of colonies in clonogenic assay, stained with crystal violet as above, were subjected to morphological analysis for quantification of area, circularity, compactness, solidity and other characteristics.

### mRNA sequencing and analysis

RNA was isolated from cells using RNeasy Plus Mini Kit (74136,QIAGEN). Raw RNA-seq expression values were normalized using DESeq2 and differentially expressed genes (log fold change level of X, and significance level of Y) from all possible pairwise comparisons (IR/C, IR/IS, IS/C) were used to visualize and cluster expression levels across the three groups: Control, IS and IR. The scripts used for downstream analysis can be found in (supplementary/github link).

### Statistical Analyses

Statistical analyses were performed with Student’s t-test or ANOVA as indicated using GraphPad Prism software. The following symbols were used to indicate significant differences: ns, *p* > 0.05; *, *p* < 0.05; **, *p* < 0.01; ***, *p* < 0.001; ****, p<0.0001.

## Results

Our previous data strongly suggested that IR adipocyte-origin exosomes would reprogram 4T1 cells, as an animal model for metastatic TNBC, to a more aggressive phenotype compared to IS adipocyte exosomes [31]. Here, we first compared the expression level of a set of genes, known to be involved in EMT, in 4T1 cells that were treated with IS or IR adipocyte exosomes *in vitro*. After 72 h, the expression level of a representative set of EMT genes increased significantly in the IR group compared to control and IS groups (**Fig. 1A**). The full dataset is available in Supplementary material (Supplemental Fig. S1D). We also observed an overall increase in EMT genes in IS compared to control, albeit less prominent than IR *vs*. control (**Fig. 1A**). We next used Ingenuity Pathway Analysis software for analysis and prediction using the gene array data and determined that several canonical pathways important for cancer progression were predicted to be upregulated, including EMT, WNT signaling and HIF1-α signaling, whereas PTEN signaling, a tumor suppression pathway, was downregulated in both IS and IR compared to control (Supplementary Fig. S1E). Notably, the senescence pathway was predicted to be elevated in the IS group compared to both control and IR. In terms of disease and function (**Fig. 1B**), the analysis predicted an elevation of progression, invasion, metastasis, invasiveness as well as angiogenesis in exosome treated groups (*i.e*., IS and IR) but a down-regulation of apoptosis and necrosis, strongly reinforcing conclusions we have already published for human breast cancer cells and human primary adipocyte exosomes [31]. To determine whether exosomes from IR adipocytes increased cell mobility, we performed a migration assay (**Fig. 1C**). Quantification showed that the IR group exhibited increased migration, whereas the IS group exhibited no change compared to control (**Fig. 1D**). Thus, these experiments showed that 4T1 cells from the IR-exosome treated group showed higher expression of EMT-related genes and increased mobility *in vitro*.

**Figure 1.**
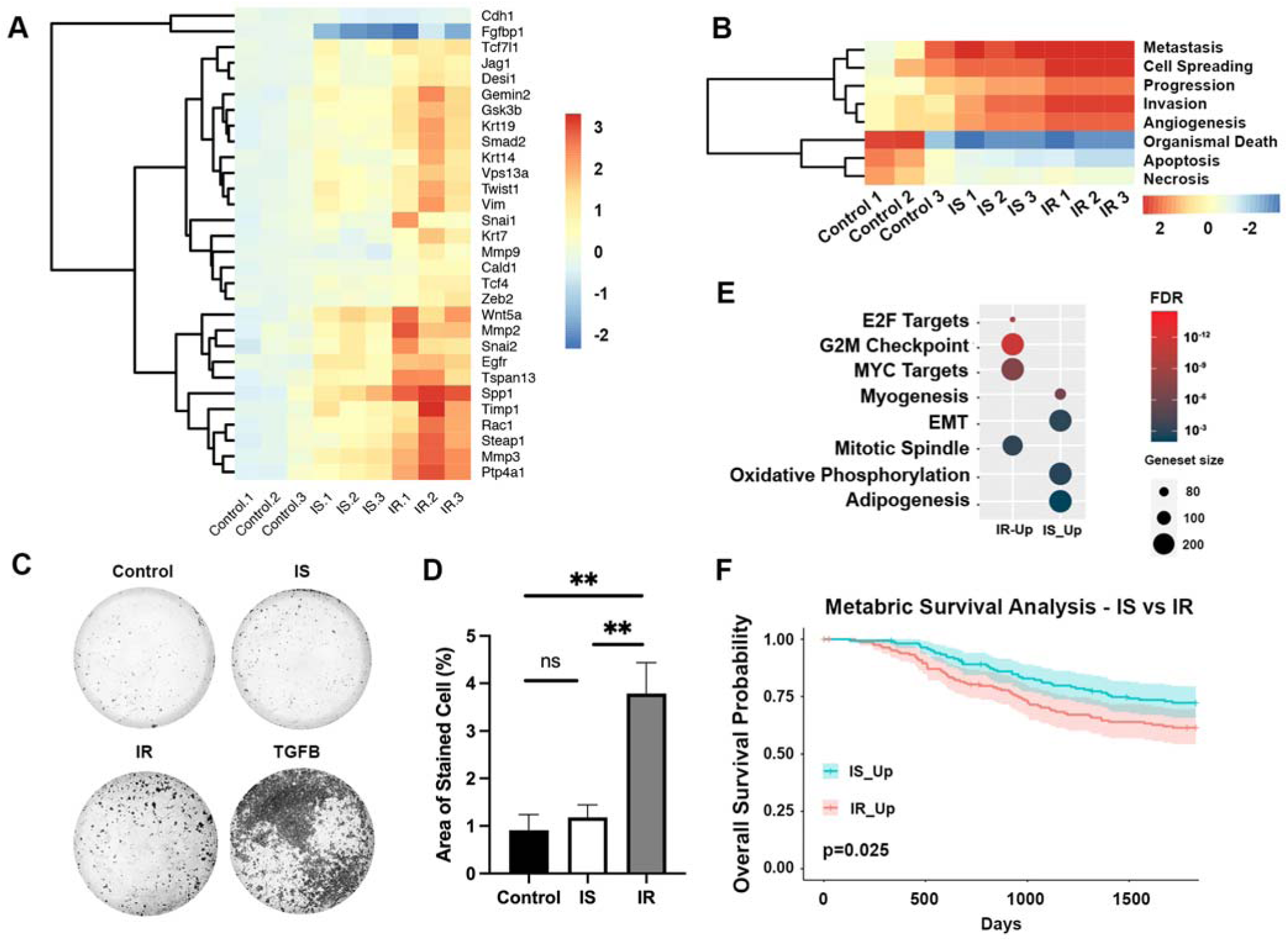
Exosomes derived from IR adipocytes increase transcription of EMT genes and migration ability of 4T1 cells *in vitro*. A. 4T1 cells were treated with IS and IR adipocyte-derived exosomes compared to control (no treatment); cellular mRNA was analyzed by EMT array. B. Ingenuity pathway analysis of differentially expressed EMT array genes. C. Representative images of migration assay of exosome-treated 4T1 cells. D. Quantification of C (n=4, *, p<0.05, **, P<0.01). E. Pathway enrichment analysis of differentially expressed genes; pathways are compared by exosome status. Bubble size indicates number of genes. Color bar indicates false discovery rate (FDR); red represents lower value, blue represents higher value. F. Survival analysis of breast cancer patients in Metabric Basal data using genes differentially expressed between IS and IR states.

In order to obtain an unbiased transcription profile, RNA-seq of 4T1 cells from the same groups was performed. Results showed a similar trend: EMT-related genes were differentially upregulated when IR was compared to IS and control (Supplementary Fig. S1F). In pathway enrichment analysis using the KS test, pathways important for cancer development were upregulated in the IR group compared to IS, including the MYC and E2F pathways (**Fig. 1E**). Pathways related to mitotic spindle function were also upregulated in IR vs IS groups, suggesting that mitosis is increased 4T1 cells treated with IR exosomes compared to IS counterparts (**Fig. 1E**). In terms of the EMT pathway, it is interesting that we only saw this pathway enriched in gene sets that are downregulated in the IS group compared to IR. We then mapped the IR vs IS differentially expressed gene sets to a METABRIC dataset and found that IR vs IS status stratified human breast cancer survival data (**Fig. 1F**), with lower survival in the IR group.

To determine whether the elevated EMT gene signatures and migration ability in the IR group translate to a physiological model, we employed the well-established 4T1 *in vivo* model for TNBC metastasis (40). 4T1 cells were treated with IS and IR adipocyte exosomes for 72h and then injected into fourth mammary fat pads of BALB/cJ mice (**Fig. 2A**). Palpable tumors formed approximately one week after injection. By the second week, primary tumors, and distant organs including the lung and brain, were harvested to quantify metastatic burden and histological changes. This model provided a valuable platform to investigate the effects of adipocyte exosomes on tumor growth and metastatic behavior.

**Figure 2.**
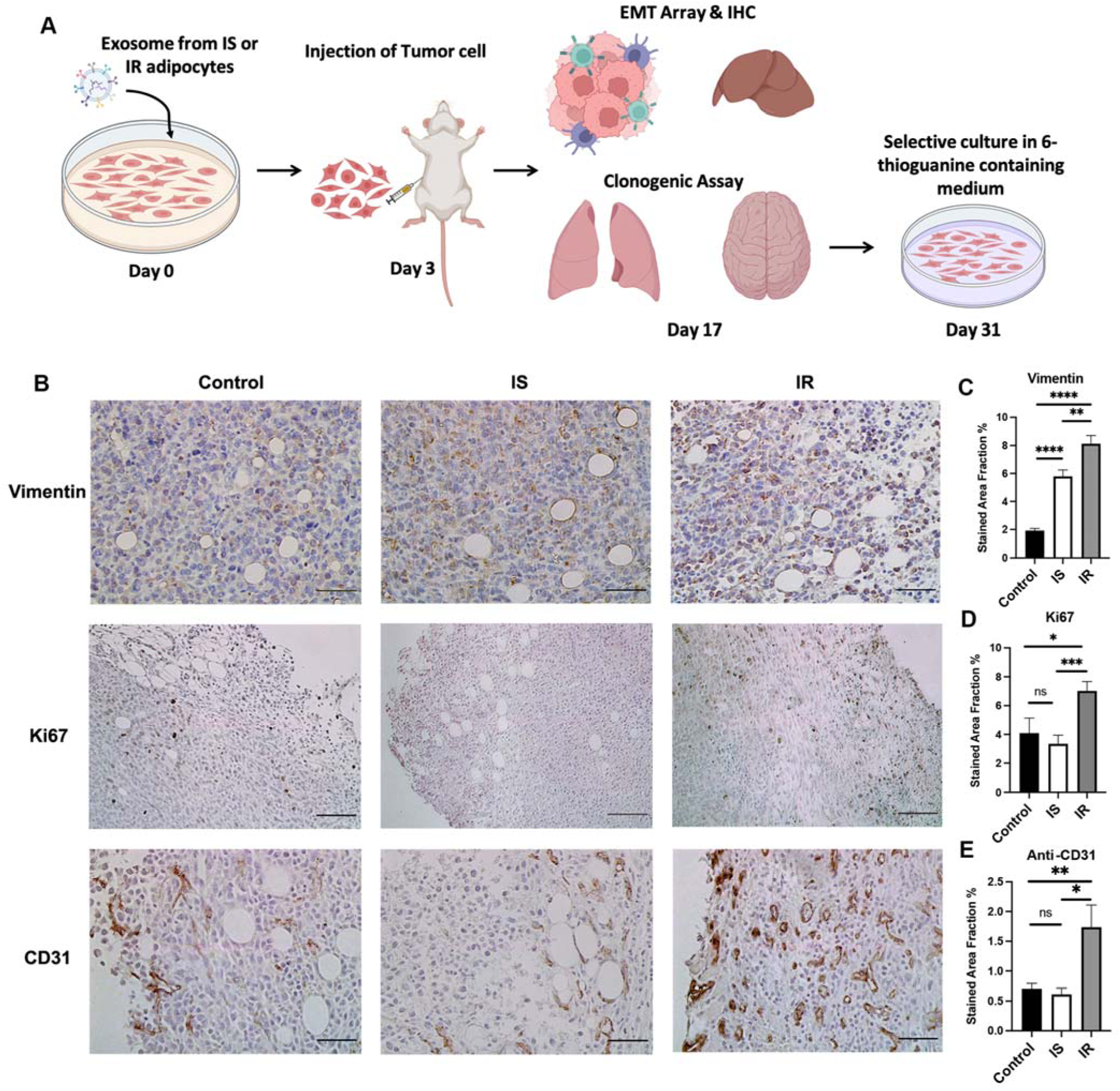
Expression of EMT, proliferation and angiogenesis marker proteins in primary tumor *in vivo*. A. Timeline of experimental procedure. B. Representative images of IHC staining with Vimentin, Ki67 and Cd31 antibodies in Control, IS and IR groups, scale bar represents 100 µm in Ki67 staining and 50 µm in Vimentin and Cd31 staining. C-E. Quantification of vimentin (C), Ki67 (D), Cd31 (E) antibody staining. (n=4, *, p<0.05, **, P<0.01)

To interrogate phenotypes in the primary tumor, we obtained FFPE sections of primary tumor and stained for Vimentin (as a marker of EMT) and Ki67 (as a marker of proliferation) (**Fig. 2B**). As expected, Vimentin and Ki67 staining were significantly higher in the IR group *vs* control and IS groups (**Fig. 2D**, **Fig. 2D**), consistent with the higher level of EMT and proliferation. Similarly, Ki67 expression in liver (Supplemental Fig. 2A), showed the same trend; Ki67 expression was higher in the IR group than IS and control. During dissections of primary tumors, we noticed markedly greater blood supply to tumors in the IR group compared to control and IS groups (*data not shown*), consistent with upregulated angiogenesis pathways we had previously reported for human breast cancer cells treated with human primary IR adipocyte exosomes [31]. Therefore, we stained primary tumors with anti-Cd31 antibody against platelet/endothelial cell adhesion molecule-1 (PECAM-1) as a vascular marker. The stained area is presented as dark brown in the image. Quantification confirmed that tumors of the IR group have a significantly higher Cd31 expression compared to control and IS groups (**Fig. 2E**). Therefore, compared to controls, exosomes derived from IR adipocytes significantly increased EMT gene expression, proliferation and angiogenesis in primary tumors of the 4T1 model of TNBC.

Beyond analysis of primary tumors, we wanted to characterize distant metastases, which are a highly informative feature of the 4T1 model. To measure metastasis in distant organs qualitatively and quantitatively, we took advantage of the unique property that 4T1 cells possess: resistance to 6-thioguanine, which enables the use of a clonogenic assay to culture the digested tissues and organs and select for metastatic clones. Only 4T1 cells survive and proliferate in 6-thioguanine containing media. We examined metastatic 4T1 clones from digested lungs after selection and found no significant difference in the stained area of the clones among any of the experimental groups (supplementary Fig. 3A). However, in brain (**Fig. 3A** top), after quantification, the stained area of the clones was significantly higher in IR than in IS and control groups. Interestingly, IS showed a significant decrease in area compared to control (**Fig. 3B**). Upon closer visual inspection of the stained colonies (**Fig. 3A** bottom) we noted differences in morphology of the cells in the colonies, with markedly more mesenchymal morphological features found in IR vs IS clones. We then measured individual cells by key parameters of cell morphology: area per cell, equivalent diameter, solidity, compactness, eccentricity, extent, and circularity (form factor). These features associate with EMT. The area per cell was smaller in the IR group than in the other two groups (**Fig. 3C**), suggesting that the similarly sized colonies in the IR group contain a greater number of mitotic cells than the other two, consistent with upregulated proliferation (**Fig. 2D**). Equivalent diameter (**Fig. 3D**) refers to the diameter of a circle that possesses the same area as the cell, which indicated the area of individual cells in IR group is significantly smaller than cells in both control and IS groups, whereas there was no significant difference observed between control and the IS group. We suggest that a higher mitotic rate of 4T1 cells in the IR group is consistent with smaller and more numerous cells than the other two groups. Solidity (**Fig. 3E**) measures minimal circumference and is generally considered a differentiator of cells with protrusion or irregular shape versus round cells. Perfectly round cells have a solidity value of 1 and cells with irregularities or holes will have a lower value, closer to 0. Cells in the IR group had solidity values significantly lower than control and IS groups, whereas there was no observed difference between control and the IS groups. Compactness is a shape descriptor used to differentiate between more compact or round objects and more elongated or irregular ones, calculated by dividing the perimeter of the object squared by its area. Cells in the IR group showed significantly elevated compactness (**Fig. 3F**), a higher perimeter/area ratio, compared to control and IS, and a more elongated and irregular shape. Like solidity, there was no significant difference between control and IS.

**Figure 3.**
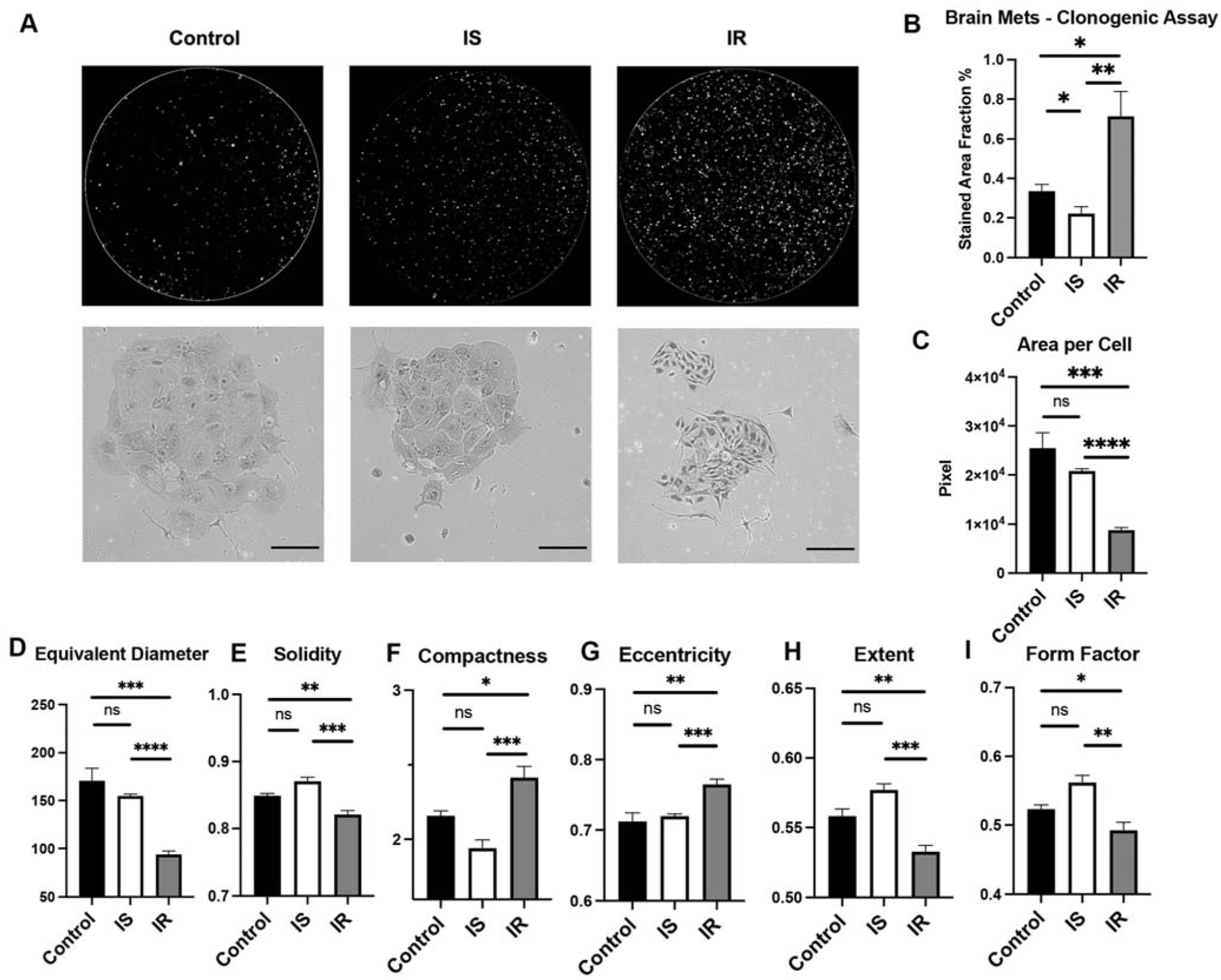
Quantification and morphology analysis of clonogenic assay from brain metastases. A. Representative images of stained colonies (top) and morphology of cells in colonies (bottom) B. Quantification of clonogenic assay from brain metastases (n=5). C-I. Quantification of area per cell (C), equivalent diameter (D), solidity (E), compactness (F), eccentricity (G), extent (H) and form factor (I) from A (images>10 per group) (*, p<0.05, **, P<0.01, ***, p<0.001, ****p<0.0001)

Eccentricity is calculated as √(1 - (b² / a²)), where ‘a’ and ‘b’ are the lengths of the semi-major and semi-minor axes of the object, respectively. For a perfectly circular object, ‘a’ and ‘b’ are equal, resulting in an eccentricity of 0. As the shape becomes more elongated, the eccentricity value approaches 1. Cells in the IR group exhibited significantly higher eccentricity compared to both the control and IS groups, while there was no statistical difference in eccentricity between the control and IS groups, indicating a more elongated cell morphology in the IR group than control and IS (**Fig. 3G**). The extent value ranges from 0 to 1. A cell with a perfect circle or ellipse shape and no protrusions would have an extent value close to 1, whereas irregularly shaped cells or cells with protrusions have lower extent values, indicating that their area deviates more from the bounding rectangle’s area. IR group had significantly lower value than control and IS, suggesting a more irregular shape and protrusions than the other two groups (**Fig. 3H**). Consistent with extent, form factor, also known as circularity, is a shape descriptor that quantifies how closely an object’s shape resembles a perfect circle, with higher values indicating a closer resemblance to a circle. Cells in the IR group exhibited significantly lower circularity compared to both the control and IS groups, indicating that their shapes deviate more from a perfect circle. However, there was no statistical difference in cell circularity between the control and IS groups, indicating that their circularity is the same (**Fig. 3I**). Taken together, these observations indicated that IR exosome treatment induces several related morphological parameters that are characteristic of mesenchymal cells, and that IS and control groups are not significantly different from each other. Thus, exosome treatment *per se* does not account for the cellular phenotypes, rather the different payloads of the exosomes contain the active factors.

Given the significant changes in morphology of 4T1 cells from brain metastases in the IR group, we expected also to see major changes in genome-wide transcription, so we undertook bulk RNA-seq analysis to investigate the whole transcriptome in a comprehensive and unbiased fashion. For each set of differentially expressed genes (**Fig. 4A**), we tested for enrichment of hallmark gene sets as well as gene sets from Qiagen using the KS score test (FDR<0.01). Significantly enriched pathways from gene sets that were upregulated in IR vs IS and IS are highlighted (**Fig. 4B**). Pathways related to mitotic spindles were enriched, indicating that proliferation was increased in the IR group compared to IS, consistent with the greater number of cells observed per colony (**Fig. 3 A-C**) and Ki67 results (**Fig. 2B**). Similarly, angiogenesis and hypoxia pathways were also enriched in IR *vs* IS, followed by EMT and cancer stem cell-related gene sets, suggesting a more mesenchymal and aggressive phenotype in the IR group. When we compared control and IS **(Fig. 4C)**, we noted that pathways that were upregulated in control included EMT, Mtorc1 and Myc targets, suggesting that some pathways in the control group have a more mesenchymal pattern than IS.

**Figure 4.**
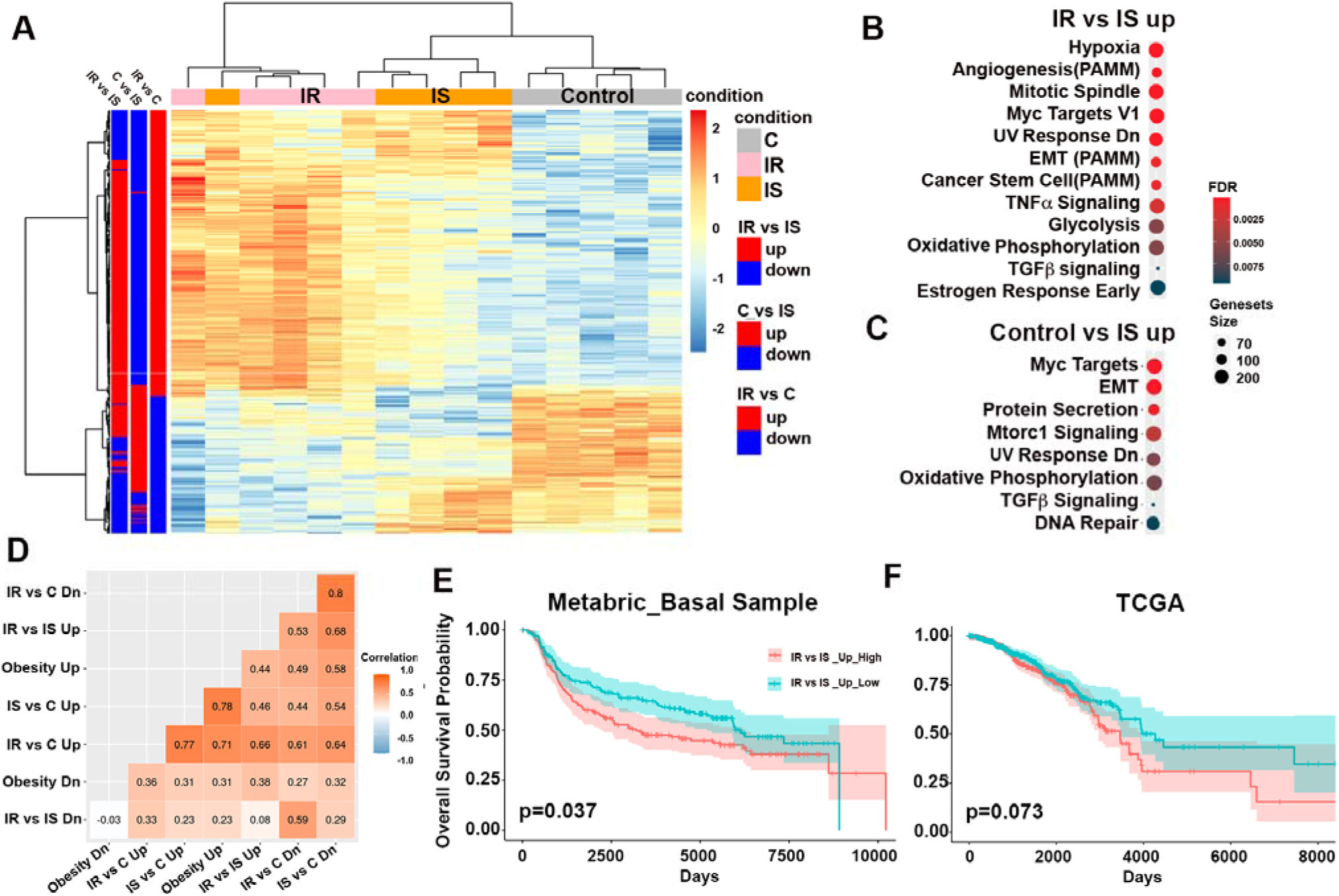
Transcriptome analysis of brain metastatic cells. **A.** Differentially expressed genes across Control, IS and IR group. **B.** Pathway enrichment analysis using genes upregulated in IR comparing to IS. **C.** Pathway enrichment analysis using genes upregulated in Control comparing to IS. **D.** Correlation matrix across all differentially expressed genesets. The row and column labels indicate the differentially expressed genesets. The Spearman correlation was calculated for each pair of results. The absolute value of these correlations is plotted in the heatmap. **E.** Survival analysis of breast cancer patients in METABRIC Basal data using differentially expressed genes between IS and IR. **F.** Survival analysis of breast cancer patients in TCGA data using differentially expressed genes between IS and IR.

To assess the clinical relevance of these patterns of gene expression, we projected publicly available breast cancer samples from METABRIC onto these gene sets using gene set variation analysis (GSVA). Using GSVA scores, we conducted a survival analysis examining the differential survival probabilities between samples expressing high ‘IR/IS UP’ scores (greater than the median) and samples expressing low ‘IR/IS UP’ scores (lower than the median). Samples expressing high ‘IR/IS UP’ scores had a lower survival probability than high ‘IS/C UP’ scores, when projecting the scores to TCGA (p-value = 0.073) and METABRIC datasets (p-value = 0.037) (**Fig. 4E&F**). TCGA and METABRIC samples were also projected onto other experimentally obtained signatures to examine correlations between these gene set scores and scores obtained by projecting the same samples to a transcriptomic obesity signature obtained from ER+ breast cancer (41) (**Fig. 4D**). For both TCGA and METABRIC ‘OB-UP’ was most correlated with ‘IR/C UP’ and ‘IS/C UP’ (> 0.5 for METABRIC, and >0.7 for TCGA).

Accumulated data pointed to differential expression of exosome payloads as the main mechanism that drives the aggressive phenotypes in the IR condition. Based on our recent study of human plasma exosome payloads that revealed major differences in miRNA profile as a function of patient metabolism (42,43), we focused on miRNA differences in the IS *vs* IR exosome composition. Here, miRNA was purified from exosomes by phenol:chloroform extraction and ethanol precipitation and repackaged into liposomes that were used to treat 4T1 cells *in vitro*. Migration assay showed a significant increase in motility in the IR group compared to both control and IS groups, although the IS group also exhibited increased migration ability compared to control (**Fig. 5A,B**). This result prompted us to identify miRNA differences among exosome types. miRNA sequencing showed differentially expressed small RNA between exosomes derived from IS and IR adipocytes (Supp Fig. 5). For quality control purposes, miRNA counts were also examined to confirm the abundance of each miRNA and biological significance. The count for miR-145a-3p was higher in IR exosomes than IS **(Fig. 5C)** and has been implicated in hypertriglyceridemia and insulin resistance (44). MiR-145a-3p also shares complimentary sequence with 3’ UTR region of *CDH1*, a mesenchymal-to-epithelial transition (MET) gene, and *PTEN*, a tumor suppressor gene, as shown by target scan analysis (**Fig. 5D**). Next, to confirm function, miR-145a-3p was obtained as recombinant purified nucleic acid and used to treat 4T1 cells *in vitro*. Because the increased brain metastasis appeared two weeks after the exosome treatment, we sought to mimic the period when cells are not in direct contact with exosomes by extending the cell culture for one extra day without treatment. **Fig. 5E** demonstrated that treatment with miR-145a-3p, which was upregulated in the IR group, showed elevated migration comparing to scrambled miRNA control.

**Figure 5.**
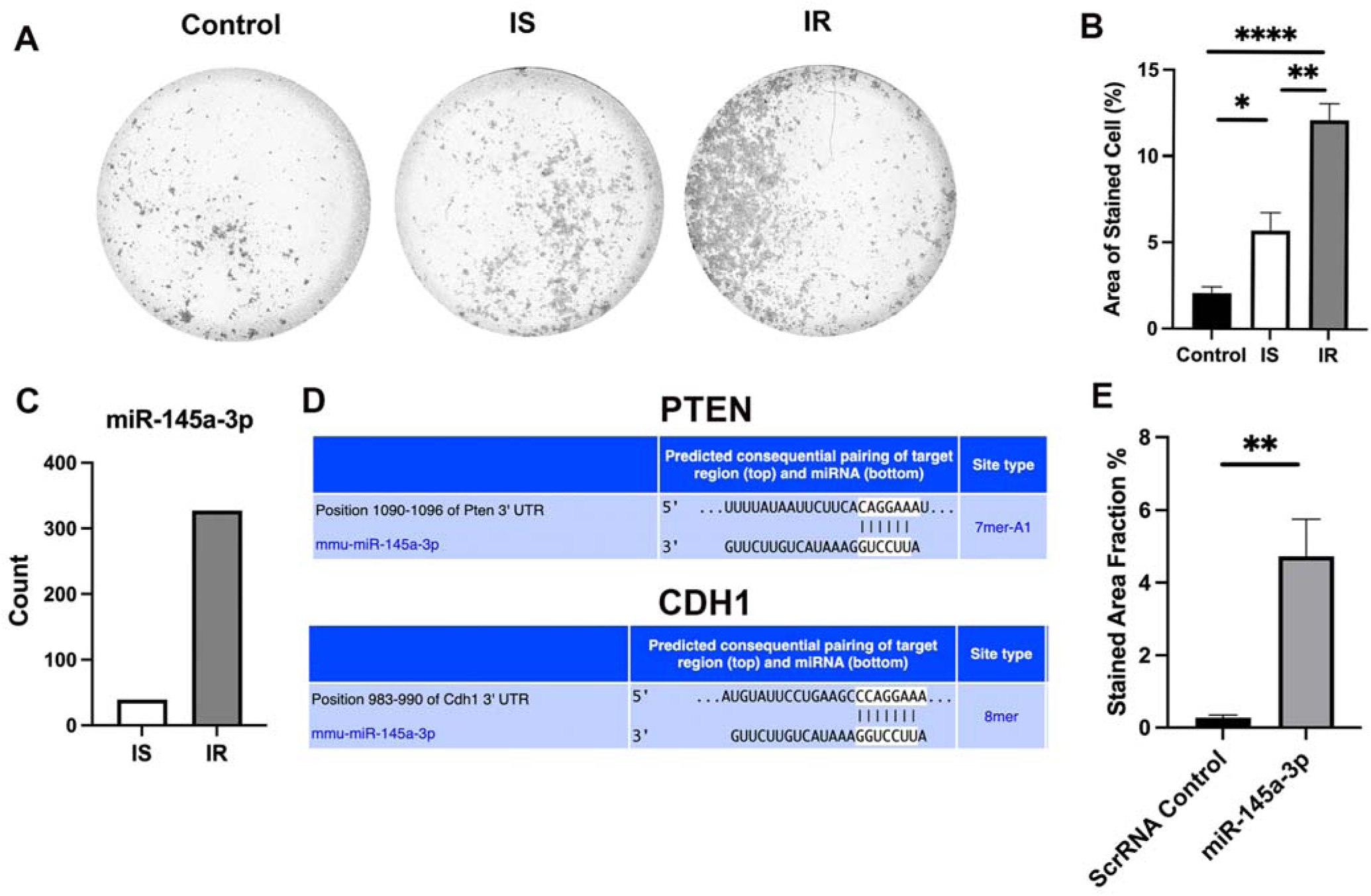
Exosomal miR-145a-3p is functional in migration assays. A. Representative images of migration assay with 4T1 cells treated with exosomal miRNA. B. Quantification of migration assay. C. miR-145a-3p is differentially expressed between IS and IR adipocyte exosomes. D. PTEN and CDH1 has complimentary sequence of miR-145a-3p in the 3’-UTR regions. E. Quantification of migration assay treated with scrambled miRNA (scrRNA) control and miR-145a-3p for 3 days and 1 day without treatment. (n=4, *, p<0.05, **, P<0.01, ***, p<0.001)

## Discussion

Population studies have established that abnormal metabolism, including IR, obesity and T2D, are risk factors for incident breast cancer [38-40]. Obesity has been positively associated with incidence of ER+ breast cancer among post-menopausal women. In addition, obesity-driven T2D has been shown in population studies to be associated with incidence, progression, and mortality of TNBC. Among breast cancer patients, TNBC has been associated with features of metabolic syndrome, particularly dyslipidemia, as likely drivers of aggressiveness. Previous research to identify mechanism and pathways of how obesity drives breast cancer has implicated several factors, including estrogen, adipokines and pro-inflammatory cytokines. However, the most critical mechanisms remain unclear in TNBC, which has the highest risks of metastasis and the fewest treatment options compared to other subtypes.

New mechanistic and potentially therapeutic possibilities have arisen with the realization that local and circulating exosomes carry molecular cargo capable or reprogramming target cells, including tumor cells, to dramatically alter transcriptional networks and cell fate in cancer. Of interest here, adipocyte exosomes [15] are newly appreciated as critical mediators of cancer progression in melanoma [17], breast cancer [16] and other tumor types [18]. We previously reported that human breast cancer cell lines dramatically upregulate gene expression networks associated with EMT, invasion, migration and cancer stem-like cell formation when challenged with exosomes purified from conditioned media of insulin resistant human adipocytes, or primary mature adipocytes differentiated from surgically isolated adipose tissue of humans with T2D, compared to IS or non-diabetic controls [32]. Here, we sought to extend these findings to *in vivo* models where metastatic behavior can be explored.

In the murine context we explore here, we note upregulation of several genes that contribute to a unique EMT signature in 4T1 cells upon induction by IR adipocyte exosomes compared to IS adipocyte exosome controls, including *Snai1*, *Snai2*, *Twist1*, *Jag1*, *Gsk3b*, *Smad2*, *Vim*, *Mmp3*, *Mmp9*, and *Zeb2*, and downregulation of *Cdh1* (**Fig. 1A**). We previously reported upregulation of the human orthologs of these same genes in the exosome-induced EMT signature we identified for the human ER+ breast cancer cell line MCF-7, where insulin resistant, adult human primary adipocytes were compared to insulin sensitive controls (32). Similar genesets were induced in MCF-7 with exosomes from primary adipocytes of adult humans diagnosed with T2D compared to non-diabetic controls (32). Only a subset of EMT-related genes is responsive to IR exosomes, and species differences likely explain additional differences between human and murine systems. Several of these genes are strongly associated with aggressiveness and metastasis of human breast cancer, including *SNAI1*, *SNAI2* and *TWIST1* (45). Furthermore, Ingenuity Pathway Analysis of gene function strongly identified invasion, metastasis and angiogenesis (**Fig. 1B**) as upregulated, with marked downregulation of necrosis and apoptosis pathways, in the murine IR exosome context compared to IS, similar to our previous report for human-only systems (32). IR exosomes likewise induced tumor cell migration in functional validation experiments, compared to controls (**Fig. 1C,D**). Projection of results onto METABRIC survival data (**Fig. 1F**) confirms the clinical relevance of these findings. We verified upregulation of EMT, angiogenesis and increased proliferation in IR exosome-treated 4T1 cells, which established primary tumors in mice, by immunohistochemistry of proteins characteristic of these processes, Vimentin, Cd31 and Ki67, respectively (**Fig. 2**), as well as in metastases to liver (**Suppl. Fig. 2A**). We conclude that exosome-driven transcriptional mechanisms are conserved across humans and mouse models and may represent a fundamental and novel mechanism regulating breast cancer proliferation, progression and metastasis.

In addition to characteristic and robust transcriptional changes associated with EMT in cell lines, IR exosomes were previously shown to induce mesenchymal morphology in human cells (32). We observed a similar phenotype *in vivo* here, including in clonally expanded colonies of brain metastases (**Fig. 3**). IR adipocyte exosomes increase metastasis in distant organs, with more mesenchymal-like cells in metastatic colonies from brain, suggesting a long-lasting change in transcriptome and genome organization. However, the increase in metastases was less obvious and statistically insignificant in lung, as 4T1 cell is highly metastatic, and lung is a vulnerable target organ even without exosome treatment. Another organ of interest is liver; where we observed higher Ki67 expression by IHC in the IR context compared to controls, however the fibrotic nature of liver tissues made clonogenic assay impossible.

Interestingly, when comparing the effects of exosomes from IS and IR adipocytes, we observed that certain gene expression changes were shared, but phenotypes did not necessarily covary. For example, both IS and IR adipocyte exosomes induced EMT gene expression, however the magnitude of the induction was greater in the IR group. Furthermore, increased EMT gene expression in the IS group did not associate with increased migration, indicating that exosome-induced EMT and migration mechanisms are independent. Similarly, in the *in vivo* model, the Cd31 and Ki67 markers for angiogenesis and proliferation, respectively, did not change in the IS group compared to control, but the EMT marker and vimentin were upregulated.

Spatially, we noticed a close association between Ki67 signal and adipocyte morphology (the open circles), as well as Cd31 and adipocytes (**Fig. 2B**), suggesting that proliferation and angiogenesis may be most active in proximity to adipocytes. This association is less obvious in the IR group compared to control and IS, with both markers more evenly distributed through the cross-section of the tumor regardless of the presence of adipocytes. Spatial assessment of the tumor microenvironment is essential to determine its exact and unbiased cell-type composition. Localization information can be crucial for determining the role each cell plays in tumor progression, and how different malignant cell subpopulations engage spatially in crosstalk with the adipocytes of the tumor microenvironment. Breast spatial tumor architecture was recently established to be directly informative to predict clinical outcome (46,47). With the emergence of spatial transcriptomic technologies, we can now also infer transcriptome-wide spatial gene expression patterns and spatial enrichment of biological processes or pathways, including those known to play a role in cancer development and progression (48). However, most spatial studies in the tumor microenvironment do not focus on adipocytes, which are often not even profiled (46,47,49). Further spatial studies are needed to explore the exosome signaling relationships among adipocytes, tumor cells and stroma in breast cancer models and patient biopsies.

The ability of 4T1 cells to survive 6-thioguanine selection enabled us to observe the morphology of the metastatic cells in the brain. The distinct differences we observed between IR and control/IS brain metastases warranted a further analysis of genome-wide transcription, especially given that the timeframe of exosome treatment is only three days. RNA sequencing results showed an increase in EMT in IR samples compared to IS (**Fig. 4A**), together with upregulation of other cancer proliferation-related pathways, such as mitotic spindle (**Fig. 4B**). Interestingly, when comparing pathways enriched in control vs IS, cancer-promoting pathways included EMT, Mtorc1, and DNA repair/proliferation were identified, indicating tumor progression may be suppressed in IS compared to control. When we examined gensets upregulated in IS vs control and IS vs IR, we found the top three enriched pathways included both interferon alpha response and interferon gamma response **(Supp Fig. 4X)**, highlighting a possibility that there was more immune response in the IS group. In addition, the G2M checkpoint pathway was enriched in IS vs control Up gene set **(Supp Fig. 4X)**, and this finding could shed some light on the tumor suppression mechanism from IS exosomes.

Mechanistically, we demonstrated that the miRNA component of exosomes contributed to the increased migration. This increase was also observed in IS group vs control, in distinction with no changes in Fig 1, implying there may be a tumor suppressive role for a protein component from IS exosomes. We have previously demonstrated that, in breast cancer cell lines, EMT gene expression and aggressiveness were increased in the IR and T2D exosome context. In this paper, we sought to support this hypothesis with *in vivo* data. The encouraging results justify the need for an observational clinical trial in which patients’ metabolism and medications are considered as modifiers of metastatic risk, especially for TNBC. Additionally, miRNAs from exosomes exhibit potential as noninvasive biomarkers to understand metastatic risk in patients with comorbid obesity and diabetes as well as therapeutic targets for the progression of TNBC.

## Acknowledgements

We thank Boston University-Boston Medical Center Cancer Center faculty R. Flynn, N. Ganem and H. Feng, for helpful comments and suggestions. We thank A. Belkina, the BUMC Flow Cytometry Core Facility, Y. Alekseyev of the Microarray and Sequencing Resource and M. Kirber of the Cellular Imaging Core Facility at BUMC for technical assistance. This work was supported by the Cancer Moonshot and Cancer Systems Biology Consortium of the National Cancer Institute (G.V.D.: U01CA182898, U01CA243004, and R01CA222170)

## Supplementary information

### Supplementary Figures and Legends

**Supplementary Figure 1.**
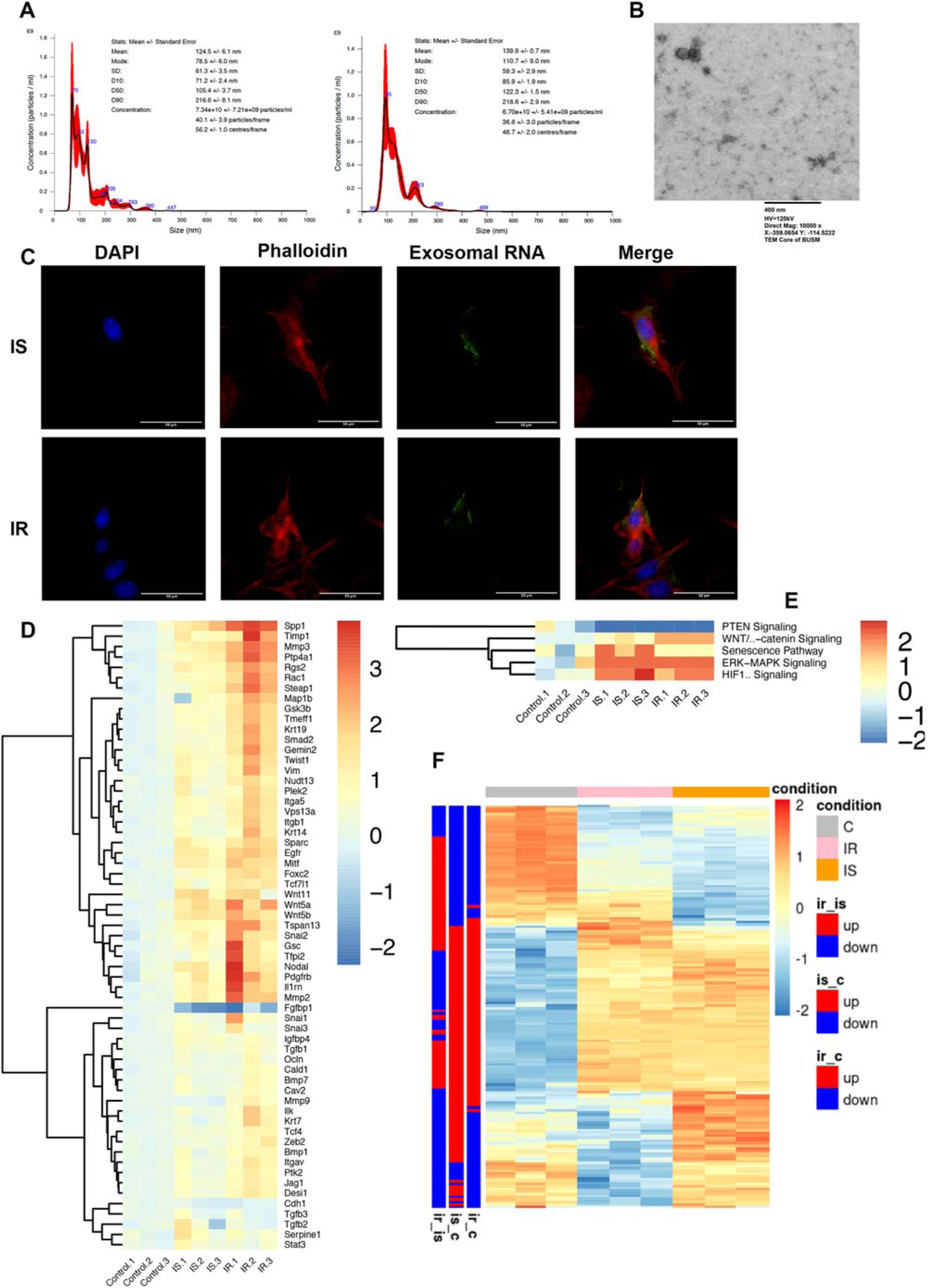
A. Histogram and statistics of exosome tracing videos (time lapse = 30s, n = 3). B. Representative image of exosomes under transmission electron microscope (TEM). C. Immunofluorescence staining of exosomal RNA intake in 4T1 cells treated with exosomes from IS and IR adipocytes. D. Heatmap of EMT gene array (60 genes) using 4T1 cells treated with exosomes for 3 days, in vitro. E. Heatmap of predicted regulation of canonical pathway via EMT gene array data. F. Heatmap of all differentially expressed genes across control, IS and IR group from bulk RNA sequencing.

**Supplementary Figure 2.**
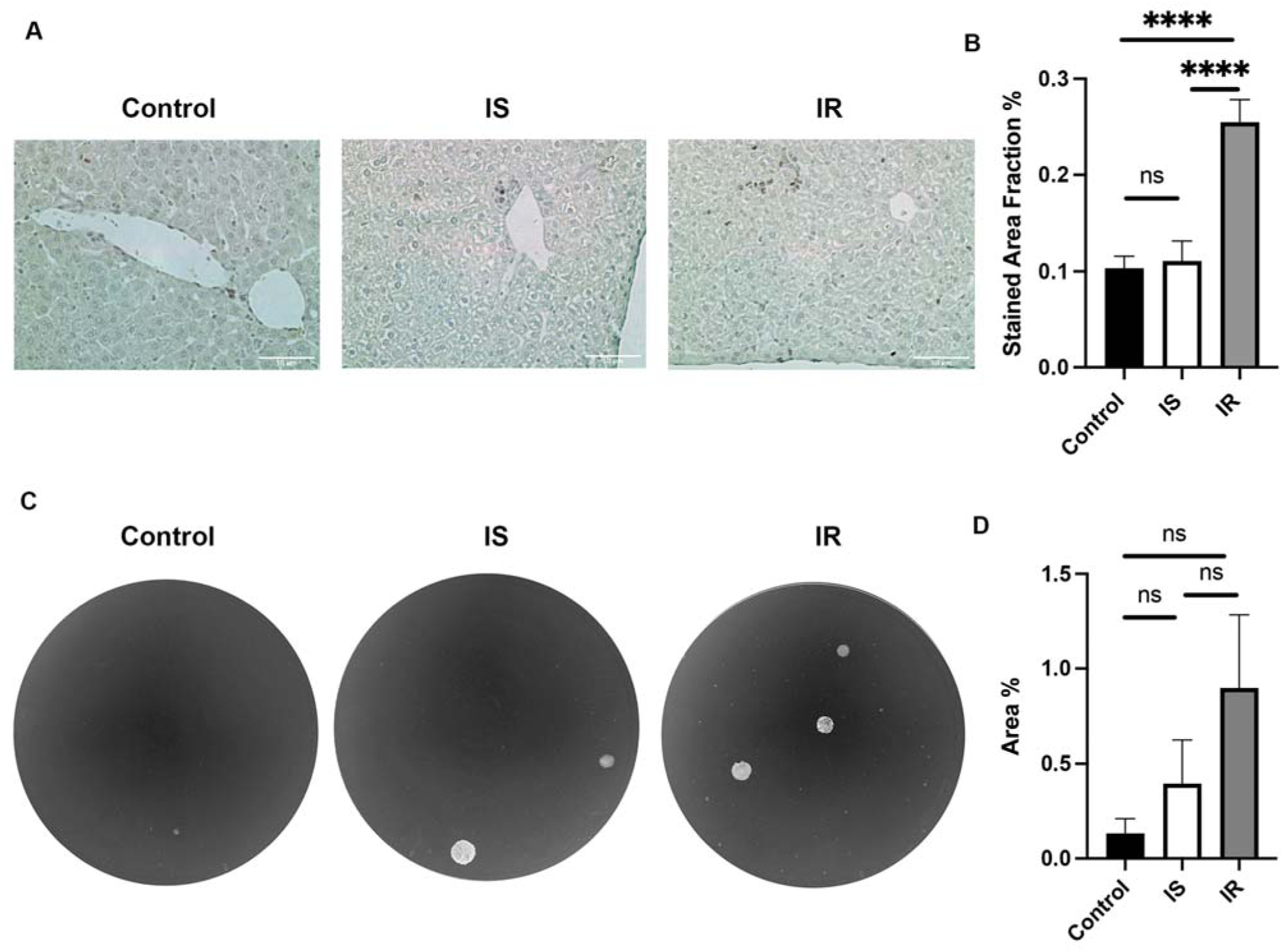
A. Representative images of IHC staining with Ki67 antibody in liver. B. Quantification of Ki67 staining (n=3, ****, P<0.0001, ns, not significant). C. Representative images of 4T1 clonogenic assay from mouse lung (n>=4, ns, not significant).

